# Rab8a-positive vesicles transport Wnt8a along cytonemes in zebrafish embryogenesis

**DOI:** 10.64898/2026.06.05.730366

**Authors:** Gemma Sutton, Lucy Brunt, Jess Bamsey, Luis Hernández-Huertas, Elizabeth Sears, Eloise Beaumont, Grace Patterson, Corin Liddle, Yang Chen, Miguel Angel Moreno-Mateos, Steffen Scholpp

## Abstract

Wnt signalling is a conserved pathway that orchestrates key developmental processes by regulating cell fate, proliferation, and tissue organisation. While the production and secretion of Wnt ligands are well characterised, less is known about how lipid-modified Wnts are delivered for long-range communication. Recently, cytonemes - actin-based signalling filopodia - have been identified as transporters of Wnts over distances to target specific cells in embryogenesis. Here, we characterise Rab8a-dependent vesicular trafficking as a crucial step in this process. Using human cell lines and zebrafish embryos, we show that Wnt ligands, such as Wnt8a, are transported with their carrier protein Wntless (Wls) in Rab8a-positive vesicles along cytonemes, where they fuse at the tips to enable hand-over and signal activation in neighbouring cells. Disruption of Rab8a function results in the intracellular retention of Wnt8a, reduces Wnt spreading and paracrine signalling, and alters embryonic patterning, consistent with reduced Wnt/β-catenin function. Conversely, Rab8a activation enhances Wnt8a dissemination, leading to increased long-range signalling and, consequently, patterning defects in embryogenesis. Our findings uncover a dedicated intracellular trafficking route for Wnt delivery to and along cytonemes, offering new insights into how the spatial precision of Wnt spreading in vertebrate tissues is achieved.

## Introduction

Wnt signalling is a fundamental pathway that plays critical roles in development and tissue homeostasis^1^. This intercellular signalling cascade, activated by secreted lipid-modified Wnt proteins, is essential for processes such as cell fate specification and proliferation^2^. The remarkable conservation of the Wnt pathway throughout the evolution of all metazoans highlights its crucial role in regulating the generation and spatial organisation of diverse cell types during development. Wnt signalling operates in both autocrine and paracrine manners, orchestrating the behaviour of ligand-producing cells and their neighbouring receiving cells.

The intracellular journey of Wnt proteins within the source cell is a complex process involving lipidation and chaperone-mediated transport to the plasma membrane^3, 4^. Following synthesis, Wnt proteins undergo critical post-translational modification, including palmitoleoylation, in the endoplasmic reticulum (ER) by the multipass-transmembrane *O*-acyltransferase Porcupine (Porcn)^5–7^. Following its displacement from the ER, the Wnt cargo receptor, Wntless (Wls)^8, 9^, transports the lipidated Wnt ligand via the Trans-Golgi Network (TGN) to the membrane for secretion.

How lipid-modified proteins with such a strong membrane affinity move in the extracellular space has remained a puzzling question; however, recent studies have highlighted the importance of specialised transport modes such as cytonemes, which are filopodia-like signalling protrusions facilitating paracrine signalling over distance during tissue patterning^4,10–12^. These actin-rich extensions can transport Wnt signalling components, including Wnt ligands and receptors, in complex tissues, enabling the precise regulation of signalling gradients in noisy and complex environments. Wnt cytonemes are dynamic structures that can extend over significant distances of about 100 µm within minutes to establish targeted contacts with receiving cells in the zebrafish gastrula, in the murine intestinal crypt, in human tumour tissue and between maturing, cortical neurons^13–18^.

While we have an increasing understanding of Wnt production and dissemination, as well as the role of cytonemes in long-range signalling, a significant knowledge gap remains regarding the transport of Wnt proteins from the production site, the ER/TGN, to the signalling site at cytoneme tips. Elucidating the mechanisms of intracytonemal Wnt transport will provide a more accurate description of how Wnt signals spread across tissues, given that extracellular transfer is likely restricted to ligand handover at cytoneme tips.

In this work, we address this knowledge gap by demonstrating that vesicles are crucial for transporting Wls together with the Wnt ligands, such as Wnt8a, in human HEPM and HEK293T cells, zebrafish fibroblasts and zebrafish embryos. We show that Wnt8a/Wls mobilisation requires Rab8a-positive vesicles travelling along the actin cytoskeleton within cytonemes. Furthermore, we provide evidence that blocking Rab8a function leads to intracellular accumulation of Wnt8a, resulting in a Wnt loss-of-function phenotype characterised by reduced paracrine Wnt target gene expression and anteriorisation of the zebrafish embryo. Conversely, the activation of Rab8a facilitates the transport of Wnt8a via cytonemes, leading to alteration of β-catenin dependent patterning of the zebrafish embryo.

By uncovering this vesicle-based trafficking route for Wnt delivery along cytonemes, we shift the view of long-range morphogen dispersal from a primarily extracellular process to one tightly regulated by intracellular Rab-mediated transport mechanisms. This reframing provides important new insights into the mechanisms that ensure precise long-range Wnt dissemination across complex tissues during development and tissue homeostasis.

## Results

### Rab8a-positive vesicles carrying a Wnt8a/Wls complex along cytonemes

Cytonemes can transport Wnt proteins to neighbouring cells, thereby orchestrating the canonical Wnt/β-catenin signalling landscape^11^. However, the intracellular mechanism of Wnt transport within cytonemes is largely unknown. To address this question, we performed a convergent screening analysis of membrane-trafficking proteins to identify vesicular markers that are present in cellular extensions and expressed early enough to control Wnt spreading during zebrafish embryogenesis.

As a discovery system for Rab-mediated endosomal trafficking, we first used human embryonic palatal mesenchyme (HEPM) cells because they model embryonic, mesenchymal cellular behaviour, including rapid proliferation and Wnt-dependent palatogenesis^19^. In a proteomic screen for cell extensions, we isolated the filopodial fraction from HEPM cells and employed an unlabelled quantitative mass spectrometry approach to compare their protein composition with that of the corresponding cell bodies. We identified a subset of Rab proteins that were significantly enriched in cellular extensions, including Rab8a, Rab13, Rab23, Rab31, and Rab35 (Fig. 1a), consistent with prior findings in other cell types^20^. Next, we hypothesised that these transporters must be fully operational during early embryogenesis - and thus maternally provided - to enable controlled Wnt signal dissemination via cytonemes and thereby control tissue patterning in the early zebrafish blastula^13,21^. Therefore, we used ribosome profiling^22,23,24^ to quantify ribosome-protected mRNA fragments of Rab proteins, and we identified 3 Rab proteins that are maternally provided and translated in the late blastula: Rab8a, Rab13, and Rab35 (Fig. 1b).

**Figure 1.**
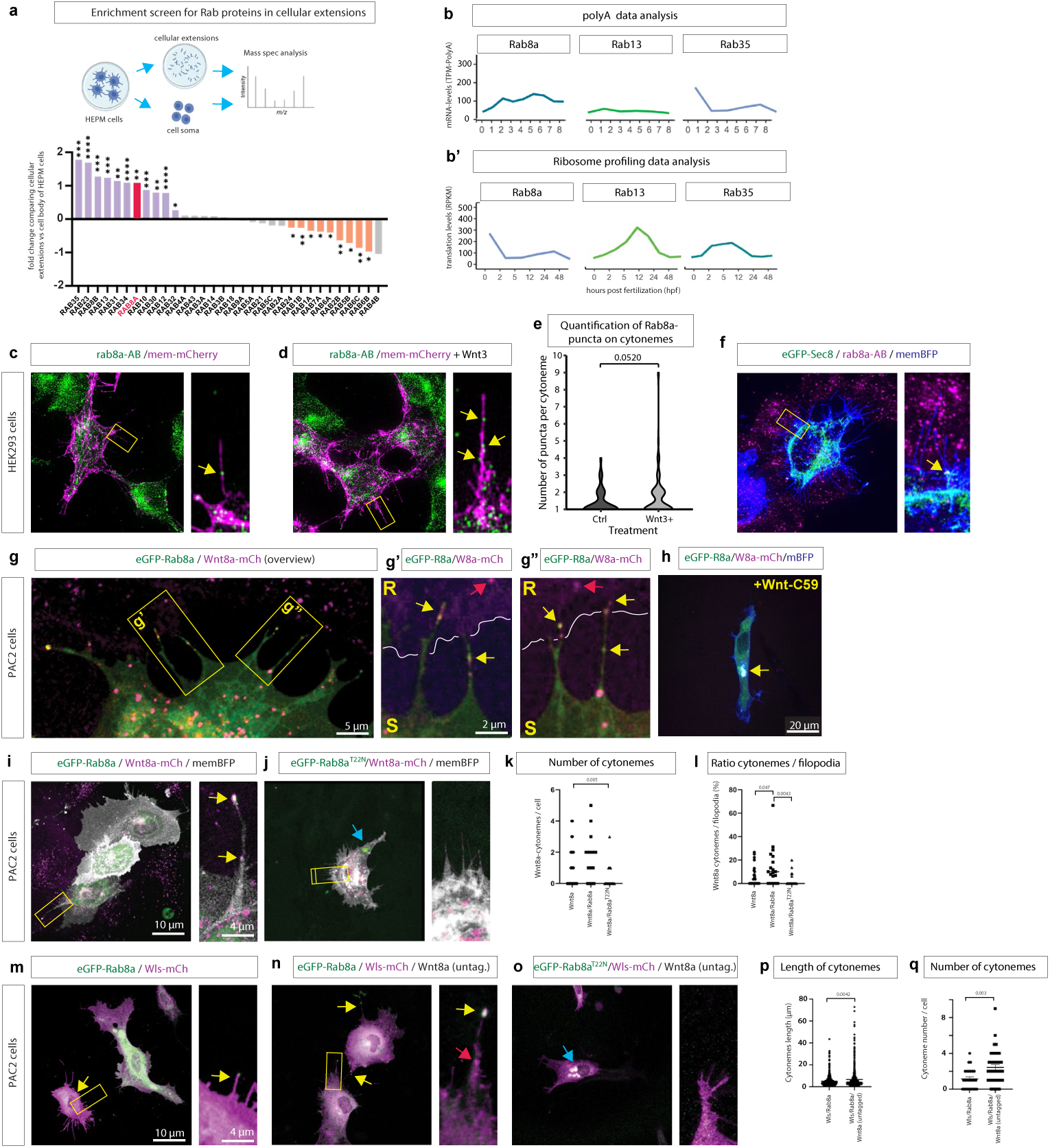
Rab8a is enriched in cytonemes and promotes cytoneme-associated Wnt8a trafficking *in vitro*. **a**, Proteomics screen quantifying enrichment of Rab GTPases in cellular protrusions versus cell bodies in embryonic cells. **b,** Poly(A RNA and ribosome profiling–based developmental expression analysis indicating maternal provision **(b**) and early translation of *rab8a* **(b’)**. **c, d,** IHC analysis of Rab8a in HEK293T cells transfected with mem-mCherry (**c**) or with the addition of Wnt3 (**d**) reveals an increase in endogenous Rab8a puncta in cellular extension after Wnt3 overexpression . Regions of interest are indicated by yellow boxes and shown enlarged in the corresponding higher-magnification images. **e,** Quantification of Rab8a puncta in cytonemes. Statistical comparison was performed using a Wilcoxon rank sum test (W=3423, p=0.052). **f,** IHC analysis of Rab8a with transfected eGFP-Sec8 and memBFP indicates co-localisation at the base of cytonemes (arrow). **g,** Confocal image of PAC2 fibroblasts co-expressing eGFP–Rab8a and Wnt8a–mCherry. **g’, g”,** High-magnification images showing co-localisation of Rab8a and Wnt8a clusters along cytonemes and at cytoneme–cell contact sites (yellow arrows, receiver “R” and sender “S” sides indicated; red arrow shows Wnt8a localisation in the receiver cell). **h,** PAC2 cells treat with a Porcupine inhibitor (10µM Wnt-C59) showing enhanced co-localisation of Wnt8a and Rab8a in the TGN (arrow);. **i,** Representative PAC2 cell expressing eGFP–Rab8a, Wnt8a–mCherry and membrane marker (memBFP), arrows indicate colocalisation of Wnt8a and Rab8a on a cytoneme; **j,** shows localisation of Wnt8a in PAC2 cells expressing the transport-defective Rab8a mutant Rab8a^T22N^. Blue arrow indicates colocalisation in the cell soma. **k, l,** Quantification of Wnt8a-positive cytonemes per cell and the ratio of Wnt8a cytonemes to total filopodia under the indicated conditions. **m–o,** Rab8a co-localises with the Wnt cargo receptor Wls on cytonemes. Representative PAC2 cells expressing eGFP–Rab8a with Wls–mCherry (± untagged Wnt8a, as indicated) (**m, n**) and eGFP-Rab8a^T22N^ and Wls-mCherry **(o)**; boxed regions are shown as enlargements. Yellow, red and blue arrows indicate Rab8a/Wls colocalisation on cytonemes, Wls clusters and colocalisation in the cell soma, respectively. **p, q** Quantification of cytoneme length and cytoneme number per cell under the indicated Wls/Wnt8a/Rab8a conditions. Scale bars, as indicated.

As Rab13 and Rab35 were not detectable in Wnt cytonemes (Extended Data 1a-c), we focused on Rab8a. Using HEK293T cells as a model to investigate signalling filopodia^13^, we first performed an immunostaining and detected puncta of Rab8a (average diameter of 0.3µm) along filopodia (Fig. 1c, controls in Extended Data 1d,e). We then tested whether Wnt expression affects the abundance of these Rab8a-positive vesicles. Overexpression of Wnt3 in HEK293T cells led to an increase in Rab8a-positive vesicles along filopodia (Fig. 1d,e). Next, we investigated the origin of these vesicles by staining cells with the post-Golgi marker Sec8. We observed co-localisation of Sec8 with Rab8a at the bases of protrusions, consistent with cargo being loaded onto Rab8a-associated vesicles at the TGN and subsequently routed towards cytonemes (Fig. 1f, Extended data 1f). We then performed an expression assay to determine whether these vesicle markers co-localise with Wnt8a, one of the earliest-expressed Wnts in zebrafish embryogenesis, which predominantly regulates the β-catenin-dependent signalling gradient in vertebrate neural plate patterning and is transported via cytonemes^13,25^. Using a transient expression approach of fluorescently tagged proteins, we observed Wnt8a forming clusters in the perinuclear region, at the cell cortex, and at the plasma membrane in PAC2 embryonic fish fibroblasts (Fig. 1g). Noteably, Wnt8a clusters co-localised with Rab8a puncta within cellular protrusions consistent with cytonemes (Fig. 1g’, g”, yellow arrows). Wnt8a signal was also detected in the neighbouring cells (red arrows). When cells were treated with the PORCN inhibitor Wnt-C59, which prevents Wnt lipidation, Wnt8a co-localises strongly with Rab8a in the TGN (yellow arrow) and transfer to neighbouring cells was blocked (Fig. 1h).

Overexpression of Rab8a with Wnt8a also significantly increased the percentage of cytonemes per cell, which contain both Wnt8a and Rab8a, suggesting Rab8a transport increases Wnt8a-positive cytonemes per cell (Fig. 1i-l). Transfection of the dominant-negative Rab8a, Rab8a^T22N^, which carries a mutation in the GTPase domain, and thus locks Rab8a in an inactive, GDP-bound state^26^, significantly reduced localisation of Wnt8a to protrusions and reduced Wnt8a-positive cytonemes per cell (Fig. 1i-l). Instead of localisation of Rab8a/Wnt8a vesicles in cytonemes and at the cell cortex, we observed that Wnt8a and Rab8a^T22N^ co-localised in the cell body (blue arrows), most likely in a TGN-associated compartment^27^, in alignment with the PORCN-inhibitor-treated cells (Fig. 1h).

Wnt ligands have been shown to be chaperoned by Wls from the ER via the TGN to the plasma membrane in *Drosophila* and human cell culture ^8,9^. Here, we show that in PAC2 fibroblasts, Wls and Rab8a co-localise on protrusions as suggested previously^43^ (Fig. 1m). The addition of Wnt8a to PAC2 cells co-expressing Wls and Rab8a leads to the formation of increased Wls clusters (Fig. 1n, red arrows) and Wls/Rab8a clusters (yellow arrows), most likely including untagged Wnt8a. Consistently, the co-expression with the dominant-negative Rab8^T22N^ blocks Wls transport towards cytonemes and retains it in a TGN-like structure (Fig. 1o). Quantification of the data sets indicates that addition of Wnt8a to the Wls/Rab8a transfection significantly increased cytoneme length and the number of cytonemes per cell (Fig. 1p, q), suggesting that Wnt8a is co-transported with Wls in Rab8a vesicles along filopodia and promotes cytoneme-based dissemination.

This analysis identified Rab8a as a maternally provided trafficking factor enriched in protrusions, positioning it to influence cytoneme-mediated Wnt trafficking and paracrine signalling at stages preceding zygotic transcription.

### 2D-CLEM analysis reveals Rab8a-positive vesicles carrying Wnt8a along cytonemes

To visualise the ultrastructural organisation of cytoneme-mediated Wnt transport, we developed a two-dimensional correlative light and electron microscopy (2D-CLEM) workflow including chemical fixation of cytonemes of PAC2 cells^29^ on the EM finder grid (Fig. 2a–c, Extended Data 2). First, we used GFP-based membrane labelling enabling us to register cytoneme–cell contacts and to trace individual protrusions from their base to distal tip (Fig. 2d). In addition to cytoneme-soma contacts^13^, we also found cytoneme–cytoneme contacts (Fig. 2d). Quantitative morphometry revealed characteristic thickening of cytoneme heads at contact sites (with an average length 1658 ± 108nm; width 357 ± 64nm) and a regular shaft (with an average length 875 ± 33nm; width 535 ± 47nm; Fig. 2e). On average, PAC2 cytonemes have a diameter of 166 ± 9nm. Vesicles were present at the cytoneme base, along the shaft, and within the enlarged terminal heads with an average diameter of 42 ± 3nm (Fig. 2d, blue arrows), suggesting a continuous trafficking route extending from the producing cell towards the contact interface.

**Figure 2.**
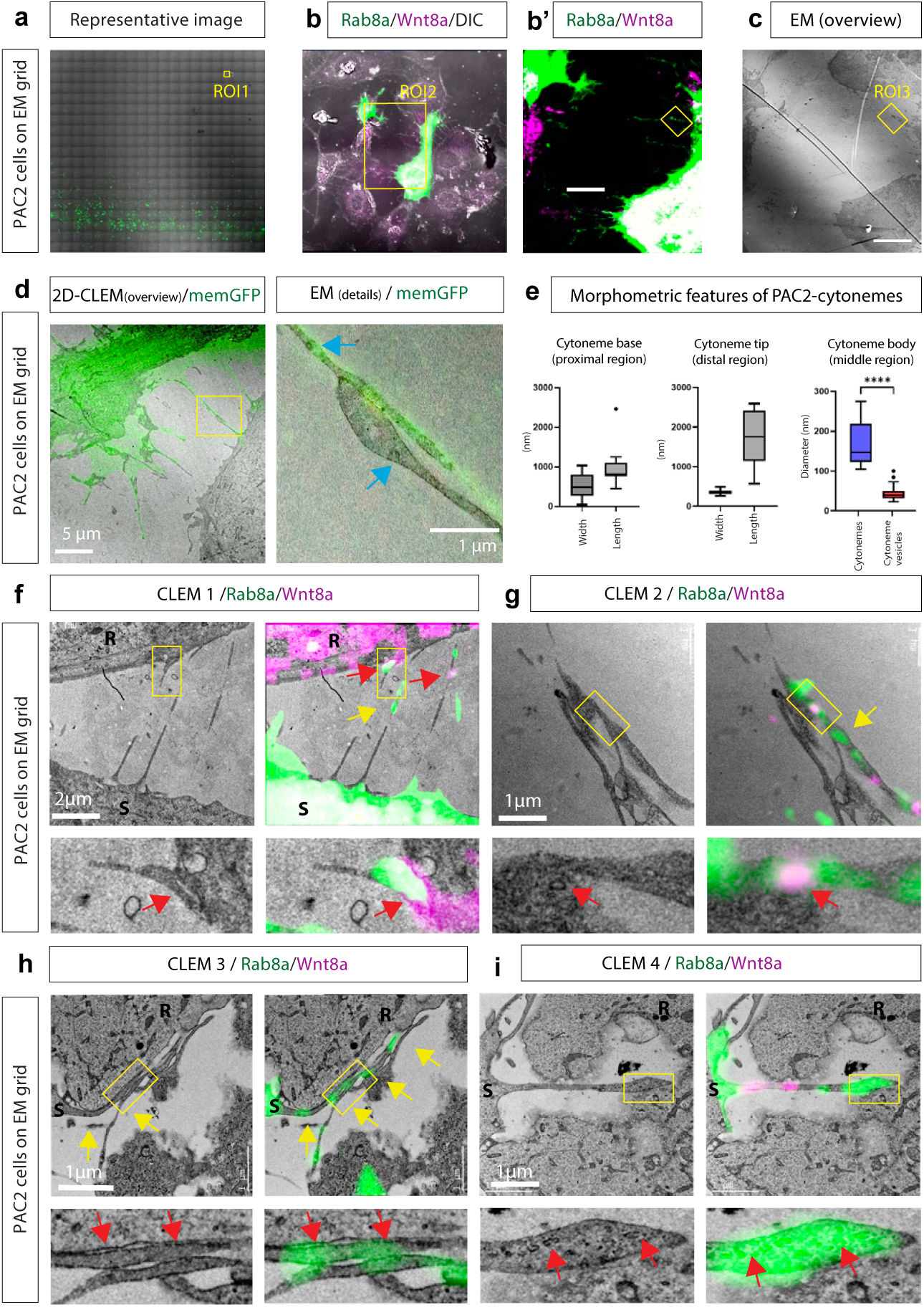
2D-CLEM reveals Rab8a-positive vesicles carrying Wnt8a along cytonemes and accumulating at specialised contact heads. **a–c,** Overview of the 2D-correlative light and electron microscopy (2D-CLEM) workflow in PAC2 fibroblasts seeded on EM finder grids: identification of transfected cells by fluorescence imaging and subsequent SEM imaging of the same regions of interest (ROIs). **d, d′,** Correlation of membrane-labelled cytonemes (memGFP) with ultrastructure in SEM; examples include cytoneme–cell and cytoneme–cytoneme contacts. Enlargements highlight vesicular structures along cytonemes (arrows). **e,** Morphometric analysis of PAC2 cytonemes quantifying terminal head dimensions versus shaft geometry and comparing cytoneme diameter to vesicle diameter (as indicated). **f–I,** CLEM in cells co-transfected with eGFP–Rab8a and Wnt8a–mCherry: fluorescence co-localisation along shafts and enrichment in cytoneme heads (red arrows) correlates with vesicular profiles in the matched EM views. Examples include Rab8a-only vesicles (yellow arrows), consistent with Rab8a carriers not being exclusive to Wnt8a cargo, and bulbous cytoneme heads densely packed with vesicles inserted into invaginations of the receiving cell surface. P, producing cell; R, receiving cell. Scale bars, as indicated.

To determine whether these vesicles correspond to Rab8a-positive transport carriers for Wnt8a, we performed CLEM experiments in PAC2 cells co-expressing eGFP-Rab8a and Wnt8a-mCherry (S, source cells) co-cultured with non-transfected PAC2 fibroblasts (R, receiver cells; Fig. 2f–i). High-resolution overlays demonstrate that Rab8a puncta co-localised with Wnt8a along cytonemes and accumulated at cytoneme tips, consistent with a role in directed morphogen delivery to neighbouring cells (Fig. 2f, red arrows). The sites of co-localisation matched the vesicular structures previously identified by EM (Fig. 2g). We also identified vesicles mapping to a Rab8a-only signal (yellow arrows), suggesting that Rab8a vesicles are not exclusive to Wnt8a cargo. At cytoneme-cell interfaces, Rab8a signal was consistently detected and often correlated with the presence of vesicles (Fig. 2h). Cytonemes could also penetrate into an invagination of the receiver cell surface, forming bulbous heads densely packed with vesicles correlating with a strong eGFP-Rab8a signal (Fig. 2i, blue arrows). In support of our previous experiments (Fig. 1g), while a Wnt8a fluorescence signal was also detected in the receiver cells, the Rab8a signal remained confined to the source cell (Fig. 2f), suggesting that only the cargo, but not the Rab8a-positive vesicular membrane, is transferred, in contrast to the transfer of Wnt5b/Ror2-positive vesicles recently reported^16^.

Our 2D CLEM approach revealed that cytonemes contain discrete vesicles along their length and expand into vesicle-rich terminal domains at contact sites. The tight apposition between cytoneme heads and the recipient plasma membrane (clefts of ∼10 nm) further implies that ligand delivery occurs through a direct, membrane-associated transfer rather than by diffusion across extracellular space. In this respect, our findings closely parallel recent observations in telocytes, where Wnt ligands are delivered to intestinal stem cells via synapse-like contacts^18^, suggesting that contact-based Wnt transfer may represent a conserved signalling mechanism.

### Rab8a-positive vesicles mediate vectorial transport of Wnt8a along cytonemes

To investigate how Rab8a-positive vesicles carrying Wnt8a navigate along cytonemes, we conducted live-cell imaging of PAC2 fibroblasts co-expressing eGFP-Rab8a and Wnt8a–mCherry (Fig. 3a). Cells were imaged for 3–5 minutes every 5 sec using high-resolution time-lapse microscopy to observe vesicle movement from the cell soma through the cytoneme to the distal tips, where ligand release occurs. (Fig. 3b).

Tracking individual vesicles showed that Rab8a/Wnt8a puncta move in a directed, sudden jumping movement along cytonemes instead of continuous motion (Fig. 3c). The average overall transport speed was 0.138 ± 0.013 µm/sec, which aligns with active transport rather than passive diffusion^30^. When stationary periods were excluded, the instantaneous speed rose to 0.232 ± 0.026 µm/sec (Fig. 3d), which falls within the range for actin-based, motor-driven cargo motion reported previously^31, 32^. Quantitative analysis of individual legs showed that vesicles typically travelled 3.7 ± 0.3 µm over 16.7 sec before pausing (Fig. 3e). These intermittent stops lasted on average 17.3 sec (Fig. 3f), suggesting transient docking or transfer events within the cytoneme. Upon reaching the distal tip, vesicles remained visible for 58 ± 11sec before disappearance, suggesting fusion or cargo release at the cytoneme head (Fig. 3g).

**Figure 3.**
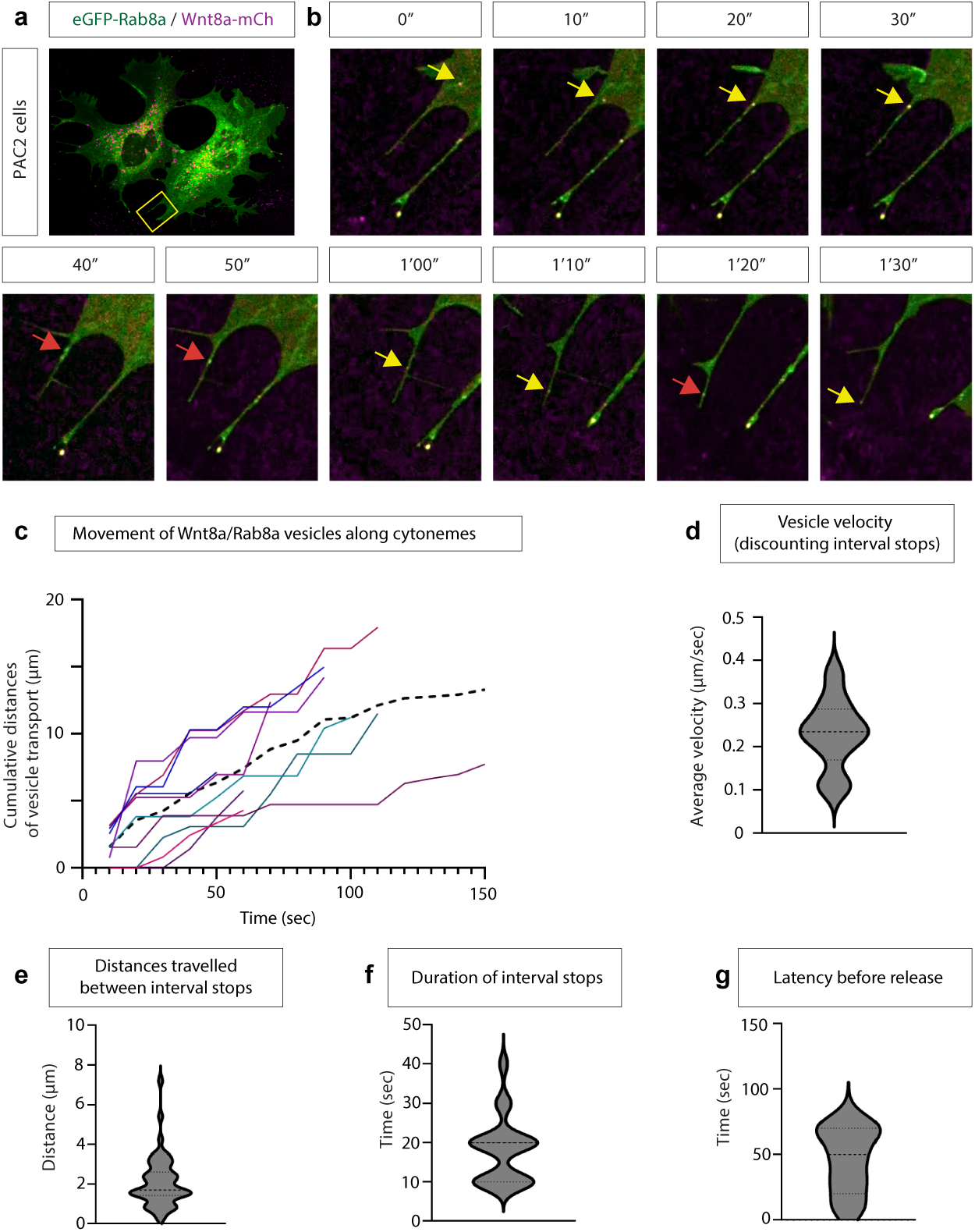
Rab8a/Wnt8a vesicles move saltatorily and directionally along cytonemes *in vitro*. **a**, Live-cell imaging setup in PAC2 fibroblasts co-expressing eGFP–Rab8a (green) and Wnt8a–mCherry (magenta). **b,** Time-lapse sequence illustrating saltatory movement of Rab8a/Wnt8a puncta along a cytoneme toward the distal tip (arrows; example frames). **c,** 10 example tracks plotted as cumulative distance travelled over time for individual Rab8a/Wnt8a vesicles along cytonemes. **d,** Distribution of instantaneous transport velocity (excluding pauses), supporting motor-driven transport dynamics along cytonemes. **e, f,** Quantification of run lengths between pauses and the duration of pausing events, consistent with intermittent stop–go movement. **g,** Dwell time of vesicles at cytoneme tips (“latency before release”), consistent with docking/fusion or cargohandover at distal heads. Scale bars, as indicated.

Together, these data indicate that Wnt8a-bearing Rab8a vesicles undergo discontinuous, motor-driven transport along cytonemes at defined velocities, consistent with active rather than diffusive movement. In combination with the CLEM analysis, this supports a model in which Rab8a-positive vesicles mediate vectorial transport of Wnt8a within cytonemes.

As Rab8a has been demonstrated to co-localise with Wnt8a on cytonemes, and the unconventional Myosin motor, Myo10, has been implicated in filopodia and Shh cytoneme dynamics^28^, we tested whether Rab8a might recruit Myo10 to drive trafficking along cytonemes. PAC2-based expression analysis of the tagged constructs showed co-localisation of Myo10 and Rab8a, and found some co-localisation of Wnt8a/Myo10 on cytonemes (Extended data 3a,b). The Wnt8a/Myo10 co-localisation could be improved by the addition of the Wnt chaperone Wls (Extended data 3c), suggesting that Myo10 can facilitate Rab8a trafficking, presumably with Wnt8a as a cargo, along PAC2 cytonemes.

### Rab8a-dependent vesicular Wnt transport along cytonemes *in vivo*

To extend our *in vitro* findings to the organismal context, we next examined the subcellular localisation of Wnt8a and Rab8a during intra-cytonemal transport in zebrafish gastrulation. We generated mosaic cell clones expressing fluorescently tagged constructs in the enveloping layer (Extended Data 4a) and analysed their localisation in living embryos at 8hpf by high-resolution confocal microscopy (Fig. 4) .

In these *in vivo* settings, Rab8a is localised on actin-positive cytonemes with an accumulation at the cytoneme tips (Fig. 4a). Wnt8a–mCherry and eGFP–Rab8a co-localised along cytonemes extending from producing cells (Fig. 4b). Within the soma of Wnt-producing cells, Wnt8a and Rab8a signals overlapped extensively, consistent with vesicular trafficking from the TGN to the plasma membrane (Fig. 4c). In contrast, Wnt8a puncta lacking Rab8a were predominantly found at the plasma membrane and in adjacent receiving cells. Notably, the Rab8a signal was almost completely absent from the receiving cells surrounding the clone, suggesting that Rab8a is recycled back to the producer cell after delivery of its Wnt8a cargo at the cytoneme–cell interface.

We next analysed whether the Wnt cargo receptor Wls uses a similar transport route. Co-expression of Wls–mCherry with eGFP–Rab8a revealed strong co-localisation in the TGN, intracellular vesicles and on cytonemes (Fig. 4d, e), further supporting a Rab8a-dependent vesicular export pathway for Wnt ligands and their transporter. The Wls signal is not detected in the receiving cells surrounding the clone, suggesting that it is also recycled back into the producing cell, similar to Rab8a. Analyses of the distance between cytonemal Wnt8a-mCherry puncta and Wls-mCherry puncta to eGFP-Rab8a puncta revealed that 68.9% and 57.6% of the puncta colocalise with Rab8a, respectively (e.g. the distance between the two puncta are <1µm) (Fig. 4f, Extended data 4b-e).

**Figure 4.**
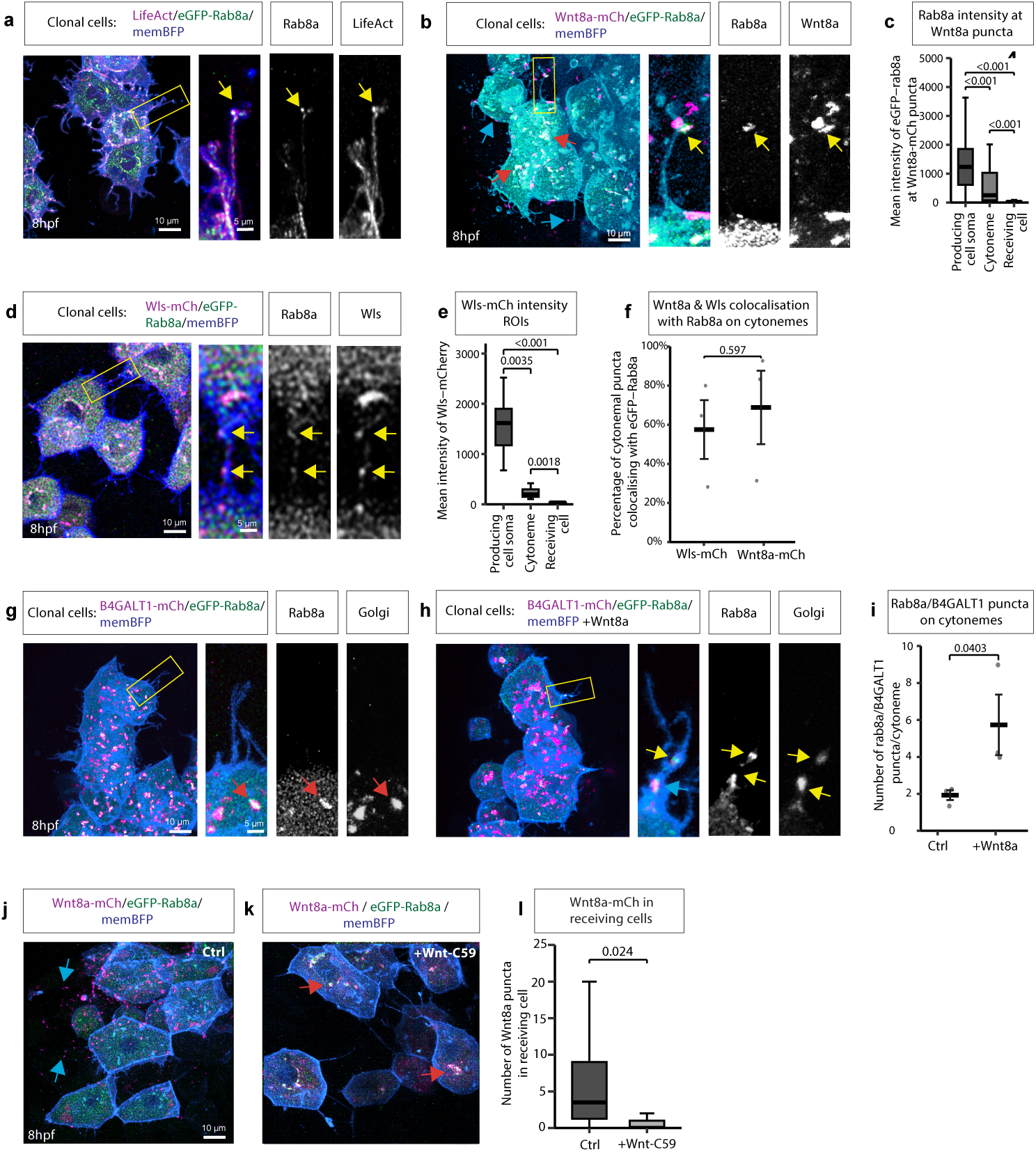
Rab8a, Wls and Myo10 co-localise with Wnt8a on cytonemes *in vivo* during zebrafish gastrulation. **a,b,d,g,h,j,k**, Representative epiblast clones in zebrafish embryos at 8 hpf co-expressing the indicated fluorescently tagged proteins, showing subcellular localisation relative to eGFP–Rab8a. The membrane marker memBFP is shown in blue. Regions of interest are outlined by yellow boxes and shown at higher magnification in the corresponding insets. Yellow arrows indicate co-localisation of the indicated markers along cytonemes, red arrows mark localisation in the cell soma, and blue arrows mark localisation in the neighbouring receiver cells. **c**, Quantification of Rab8a fluorescence intensity in vesicles co-localising with Wnt8a–mCherry. Differences in Rab8a intensity at Wnt8a-mCherry puncta in the indicated structures were assessed using a Kruskal-Wallis test, which revealed a significant overall effect (Χ^2^=233.93, df=2, p<0.001). Dunn’s posthoc test adjusted with the Holm method was used for pairwise comparison. **e**, Quantification of Wls–mCherry fluorescence intensity in the indicated subcellular regions. Statistical differences in Wls intensity in the indicated subcellular structures was assessed using a Kruskal Wallis test, revealing a significant overall effect (Χ^2^=233.93, df=2, p<0.001). Dunn’s post hoc test adjusted with the Holm method was used for pairwise comparison. **f,** Quantification of the co-localisation of Wls or Wnt8a with Rab8a-positive vesicles. Co-localisation between cytonemal Rab8a puncta and Wnt8a or Wls puncta was quantified as a binary outcome (colocalised = 1, non-colocalised = 0). A binomial generalised linear mixed-effects model (logit link) was used to test for a difference in colocalisation probability of Wnt8a and Rab8a vs. Wls and Rab8a on cytonemes (β=0.588 ±0.986, p=0.551) with biological replicate included as a random effect. **i,** Quantification of the number of Rab8a/B4GALT1 vesicles on cytonemes following Wnt8a overexpression. Welch two-sample t-test revealed that addition of Wnt8a significantly increased vesicle numbers per cytoneme (t=-2.15, df=18.82, p=0.045). Quantification of Wnt8a puncta in receiving cells after Porcupine inhibition. Poisson generalised linear mixed-effects model, with biological replicate included as a random effect, revealed a significant reduction in the number of Wnt8a puncta on cytonemes following Wnt-C59 treatment (β=-2.09 ±0.91, p=0.022). Scale bars, as indicated.

To investigate the origin of Rab8a-positive vesicles *in vivo*, we co-stained Rab8a-expressing cells with the Golgi marker, mCherry-tagged B4GALT1^33^. Rab8a strongly colocalises with B4GALT1in the producing cell soma, whereas Rab8a-positive vesicles were rarely detected within cytonemes (Fig. 4g, i). To stimulate Rab8a transport, we co-expressed untagged Wnt8a, which resulted in the appearance of Rab8a puncta within cytonemes, some of which still co-localised with the mCherry- B4GALT1 (Fig. 4h, i).

Next, we tested whether Porcupine-mediated Wnt palmitoleation^7^ is required for loading of Wnt into Rab8a-positive vesicles *in vivo*. Blocking Wnt lipidation significantly reduced Wnt8a–Rab8a co-localisation within cytonemes and strongly diminished Wnt delivery to adjacent cells (Fig. 4j-l). Furthermore, lipidation inhibition caused pronounced accumulation and co-localisation of Wnt8a with Rab8a at the TGN (Fig. 4k, yellow arrows), consistent with palmitoleation being required for Golgi exit and Rab8a-mediated cytoneme trafficking as observed previously in PAC2 cells (Fig. 1h).

Finally, we asked whether Rab8a recruits Myo10 to support Wnt8a cytoneme trafficking *in vivo* (Extended Data 4f-i). Although Myo10 was broadly present on filopodia, Wnt8a showed only a partial overlap with Myo10 (Extended data 4g). Co-expression of untagged-Rab8a increased Wnt8a–Myo10 co-localisation on cytonemes (Extended data 4h). Consistent with our analysis in PAC2 cells, the Wnt chaperone, Wls, and Myo10 also colocalise on cytonemes (Extended data 4i, 3c) This data is consistent with a potential cooperative Rab8a–Myo10 function in cytoneme-based Wnt8a delivery, although Myo10 likely has roles in cytoneme transport of other cargo.

### Rab8a function is required for Wnt8a transport in cytonemes

To probe the functional requirement of Rab8a during early embryogenesis *in vivo*, we established a transient knockdown system targeting maternal and zygotic mRNAs using RfxCas13d-mediated mRNA degradation^34–36^. Knockdown of *rab8a* caused epiboly defects, specifically developmental arrest at mid-gastrula stages (Fig. 5a, Extended Data 5). To confirm specificity, we performed a rescue assay by co-injecting *rab8a* mRNA in which the endogenous 5′ and 3′ UTRs were replaced by β-globin UTRs, rendering the transcript insensitive to Cas13d-gRNA targeting (Fig. 5b). First, we observed that gRNAs targeting the 3’UTR of *rab8a* mRNA recapitulated the developmental phenotype and that the overexpression of *rab8a* mRNA did not cause any phenotypic alteration during early development (Extended Data 5). The provision of this synthetic mRNA rescued normal development in over 50% of embryos. Then, and to further verify that the observed developmental phenotype was not caused by toxicity from the *in vitro* transcribed (IVT) products or collateral activity of the CRISPR-RfxCas13d system^36^, total RNA and ribosomal integrity were analysed in both control and knockdown samples, and comparable results were observed (Extended Data 5). Altogether, these results indicate that Rab8a is essential in early embryogenesis, most likely as a cargo transporter, and further confirm the specificity and efficiency of the maternal-zygotic knockdown approach depleting *rab8a* mRNA during early embryogenesis.

**Figure 5.**
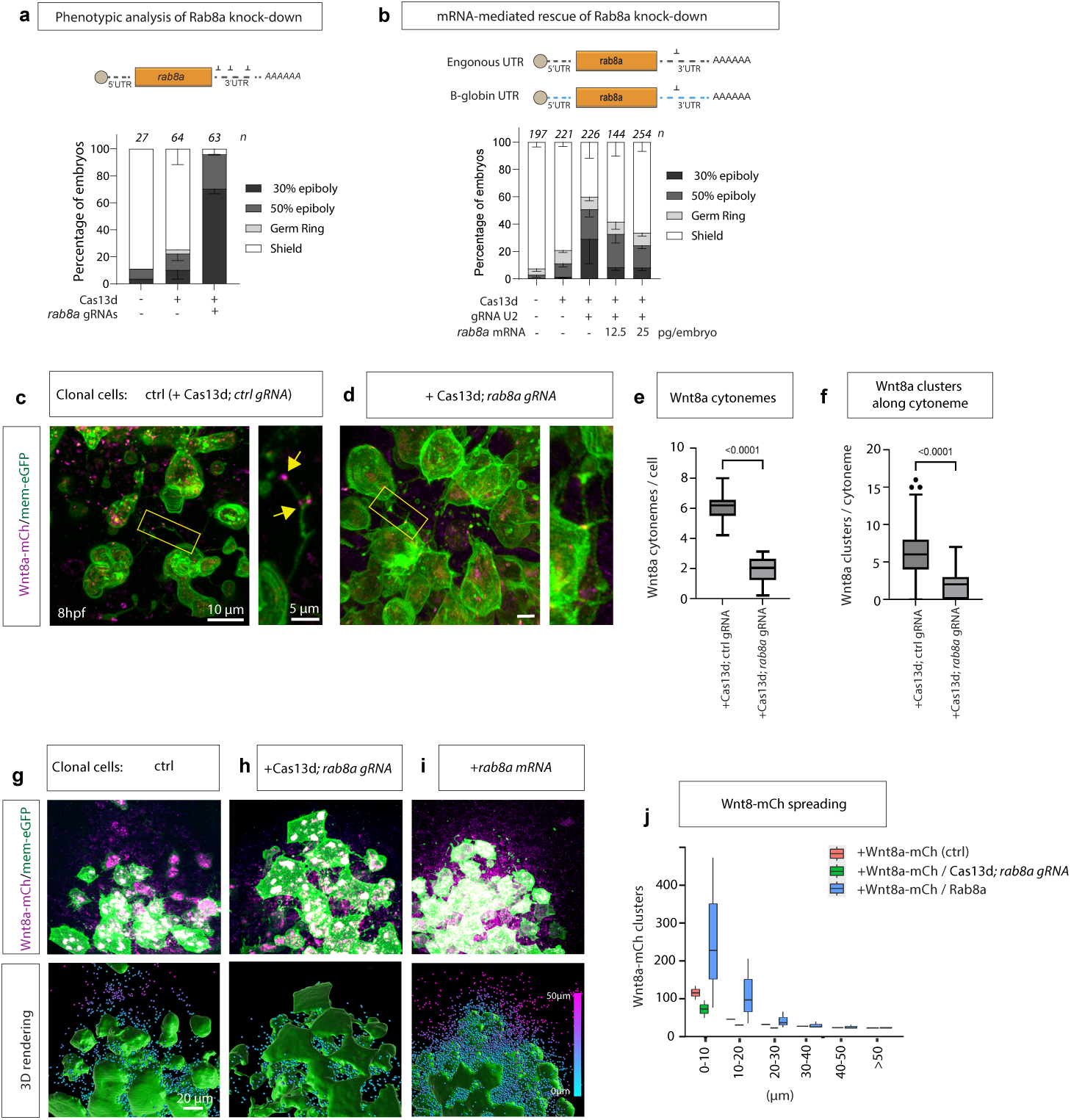
Rab8a is required for cytoneme-based Wnt8a transport and paracrine signalling *in vivo*. **a**, Maternal-zygotic knockdown strategy using Cas13d and *rab8a* gRNAs; stacked bars show developmental delay/arrest distribution across scored embryos under the indicated injection conditions. **b,** Rescue strategy using *rab8a* mRNA with β-globin UTRs to bypass gRNA targeting; dose series quantification showing partial restoration of development (bars; n as indicated). **c, d,** Mosaic epiblast clones expressing Wnt8a–mCherry together with Cas13d and either ctrl gRNA **c**, or *rab8a* gRNA (**d**; membrane marker shown in green); magnified insets highlight cytoneme-associated Wnt8a clusters (arrows). **e, f,** Quantification of Wnt8a clusters on cytonemes and frequency of Wnt8a puncta per cell/cytoneme in control versus *rab8a* knockdown conditions. **g–i,** Epiblast clones co-expressing Wnt8a–mCherry and mem-GFP, together with the indicated constructs, were imaged at 8 hpf, and three-dimensional reconstructions were generated to quantify Wnt8a–mCherry clusters in neighbouring cells. **j,** Cluster distribution was quantified as a function of distance from the source cells.

We then used this approach to assess whether Rab8a-depleted cells can still transport Wnt8a along cytonemes. To exclude the global developmental defects observed above, we generated small cell clones expressing Wnt8a–mCherry together with RfxCas13d and *rab8a* gRNA in gastrulating embryos (Fig. 5c). Compared to control clones, Rab8a-deficient cells displayed a significant reduction in the number of Wnt8a clusters along cytonemes (Fig. 5d, e) and fewer Wnt8a puncta per individual cytoneme (Fig. 5f). Next, we quantified the spatial spread of Wnt8a in the tissue surrounding these clonal cells. To this end, the fluorescence signal of Wnt8a–mCherry was measured in neighbouring receiving tissue and the tissue volume was calculated. Co-expression of Rab8a and Wnt8a in clonal cells resulted in a significant increase in the volume of Wnt8a–mCherry signal compared with clones expressing Wnt8a alone (Fig. 5g, h, i). Conversely, RfxCas13d-mediated Rab8a knockdown led to a pronounced reduction in the Wnt8a–mCherry–positive halo surrounding the clone (Fig. 5i, j).

Together, these *in vivo* experiments demonstrate that Rab8a function is essential for cytoneme-based transport of Wnt8a across tissues during zebrafish gastrulation. This supports a conserved model in which Rab8a orchestrates the vectorial delivery of Wnt ligands to target cells via vesicular trafficking rather than diffusion through extracellular space, indicating that Rab8a activity increases the effective signalling range of the ligand *in vivo*.

### Rab8a is required for paracrine Wnt8a signalling in zebrafish development

To test the consequences of altered Rab8a levels on paracrine Wnt signalling, we generated small clones expressing Wnt8a in zebrafish embryos and compared them to control clones (Fig. 6a-e). At 8 hpf, we fixed the embryos and stained them for the canonical Wnt target gene and hindbrain marker *gbx1*^37^. The expression of *gbx1* in zebrafish gastrulation depends on cytoneme-mediated Wnt8a dissemination^14^. Wnt8a-positive clones generate a halo of *gbx1*-positive cells around the clonal cells (Fig. 6b) compared to the control mRNA-injected clones (Fig. 6a). We then co-expressed mRNAs of *wnt8a* and *rab8a* in the clonal cells and observed an increased area of *gbx1*-positive cells around the clone (Fig. 6c). Conversely, when Rab8a function was blocked by the Cas13d knockdown approach in the clonal cells, we found a decrease in *gbx1*-expressing cells surrounding the clone (Fig. 6d). Interestingly, high-magnification analysis revealed that clonal, Rab8a-deficient cells expressing Wnt8a can still activate ectopic *gbx1* expression by autocrine signalling (inset, yellow arrows). In summary, quantification of the observation revealed a significant increase in the *gbx1*-positive area around the clones upon expression of Wnt8a/Rab8a, whereas a reduction of the *gbx1* expression domain was observed if Rab8a function was knocked down (Fig. 6e), suggesting that Rab8a function regulates paracrine Wnt signalling. We conclude that co-expression of Rab8a and Wnt8a enhanced induction of the posterior neural marker *gbx1*, linking Rab8a-dependent trafficking to tissue-level patterning outcomes.

**Figure 6.**
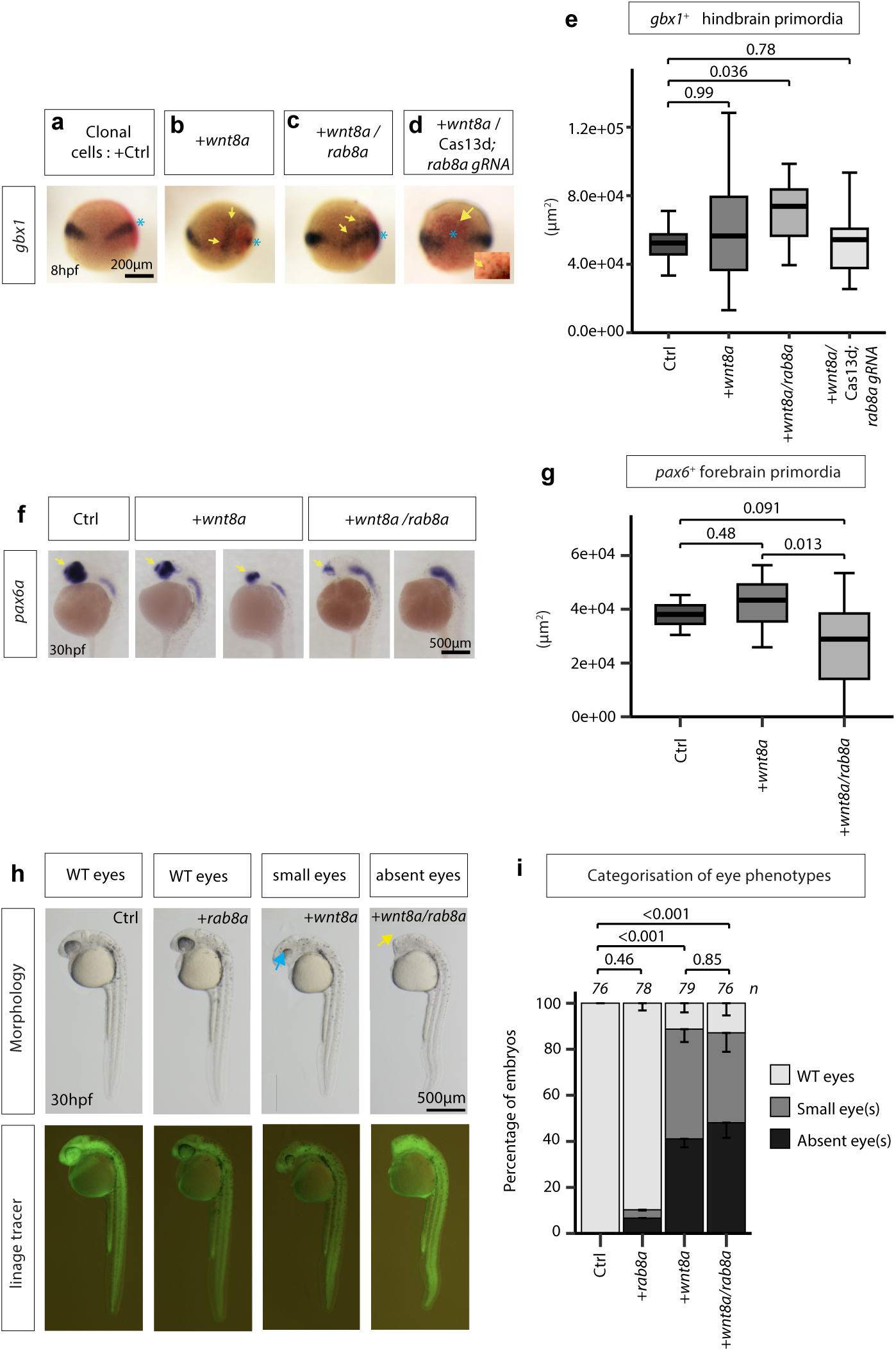
Rab8a potentiates Wnt8a-dependent forebrain patterning phenotypes. **a–d**, Whole-mount *in situ* hybridisation for *gbx1*: clones (control, *wnt8a*, *wnt8a + rab8a*, and *wnt8a* in a Rab8a knockdown background) induce *gbx1* expression (yellow arrows). Animal pole oriented to the top. **e,** Quantification of the induced *gbx1*^+^ area around clones. **f,** Whole-mount *in situ* hybridisation for *pax6a* at 30hpf in larvae microinjected at 2-4 cell stage with control mRNA, *wnt8a*, and *wnt8a*+*rab8a. wnt8a* overexpression and*wnt8a+rab8a* co-overexpression show a reduced forebrain domain, indicating increased Wnt/β-catenin signalling. **g,** Quantification of *paxCa*^+^ forebrain area (µm²) under the indicated conditions. Kruskal-Wallis test indicated a significant reduction of the forebrain area (Χ^2^=8.862, df=2, p=0.012). Dunn’s post hoc test adjusted with the Holm method was used for pairwise comparison. **h,** Representative morphology images at 30 hpf showing eye phenotypes of larvae microinjected at 2-4 cell stage with *rab8a, wnt8a* and *wnt8a/rab8a* compared to control mRNA injections. The injected cells were labelled with Dextran-Fluorescein 10,000MW (Mini-Emerald). In *wnt8a* and *wnt8a/rab8a* overexpression, eye defects were observed, including smaller eyes (blue arrows) and absent eyes (yellow arrows) **i,** Quantification of eye defects across conditions. Kruskal-Wallis test indicated a significant effect of mRNA microinjection on the eye phenotype severity (Χ^2^=212.34, df=3, p<0.001). Dunn’s post hoc test adjusted with the Holm method was used for pairwise comparison. Scale bars represent standard error.

To further assess the impact of Rab8a on anteroposterior patterning in the zebrafish, we overexpressed Wnt8a in the whole embryo via microinjection at the 2-4-cell stage, with simultaneous overexpression of Rab8a. At 30 hpf, we analysed the expression pattern of the homeobox transcription factor *pax6a*, which marks the forebrain and hindbrain regions and can, therefore, be used to visualise anteroposterior patterning in the zebrafish brain anlage. We found that overexpression of low levels of *wnt8a* mRNA (10 pg/embryo) reduced the *paxC*-positive forebrain area (Fig. 6f, g), consistent with the well-described posteriorisation function of canonical Wnt signalling in embryogenesis ^38^. Co-expression of *wnt8a* and *rab8a* mRNA led to a synergistic phenotype and a significant reduction of the *paxC*-positive forebrain area (Fig. 6f, g).

Finally, we examined the morphological phenotype of the larvae. Consistent with a reduction in *paxCa* expression in the forebrain, we found that Wnt8a- and Wnt8a/Rab8a-expressing larvae developed eye abnormalities, including complete absence of eyes or smaller eyes compared to microinjected controls (Fig. 6h,i, Extended Data 5a,b). We observe an increase in severity of the eye phenotype in Wnt8a/Rab8a-expressing larvae compared to Wnt8a overexpression, suggesting that paracrine Wnt8a signalling is promoted by Rab8a function.

## Discussion

### Importance

Wnt signalling is a fundamental organiser of embryogenesis, stem-cell maintenance, and tissue morphogenesis, and its dysregulation underlies major human diseases, including congenital malformations and cancer^1^. Despite decades of work, a central question remains unresolved: how Wnt ligands spread between cells and across tissues. As Wnt proteins are lipid-modified and membrane-associated molecules, a major focus of research has been on characterising extracellular carriers, including lipoproteins, exosomes, and soluble Wnt-binding proteins, to enable long-range extracellular spreading and, at the same time, to explain how Wnts establish a controlled signalling gradient, including specificity and robustness *in vivo*.

Our findings are consistent with an alternative mechanism in which Wnt ligands, particularly Wnt8a, remain predominantly within cells during transport. We demonstrate that transport vesicles rapidly transport Wnt8a along slender, long cytonemes across tissues. At specialised cytoneme-cell contact sites, Wnt8a is handed over to specific target cells, minimising its presence in the extracellular space significantly.

### Rab proteins control intercellular cargo distribution

Intracellular membrane trafficking is controlled by Rab GTPases, a large family of small GTP-binding proteins that regulate vesicle formation, transport, tethering, and fusion between distinct subcellular compartments^39^. To identify trafficking regulators involved in cytoneme-mediated Wnt signalling during early zebrafish development, we performed a subtraction screen for Rab proteins that are enriched in cellular protrusions (Fig. 1a) and maternally supplied as Wnt signalling is active prior to the mid-blastula transition (Fig. 1b). Our analysis identified Rab8a as a maternally provided Rab GTPase that is strongly enriched in protrusions.

Rab8 was originally described as a regulator of anterograde transport of newly synthesised proteins to the plasma membrane in MDCK cells^40^. Consistent with this function, Rab8 depletion impairs delivery of the transferrin receptor to protrusions, whereas Rab8 activation promotes transport of the vesicular stomatitis virus glycoprotein (VSV-G) to protrusive membranes in mammalian cell culture^41^. Recent proteomics analyses further show that Rab8a co-purifies with exocytic and plasma membrane–targeting proteins, including syntaxins such as SNAP23, and annexins, supporting its role in post-Golgi trafficking and polarised secretion^40,42^.

Together, these data position Rab8 as a key mediator of directed membrane and cargo delivery to protrusive cell surfaces, consistent with a role in cytoneme-associated Wnt transport during early embryogenesis.

### Link between Rab8 and Wnt

Rab GTPases regulate endosomal trafficking routes that can shape intracellular Wnt transport and thereby couple ligand handling to signalling output^3^. Specifically, Rab8a has been suggested to regulate the trafficking of Wntless (Wls) - the essential Wnt cargo receptor - from the TGN to the plasma membrane, thereby controlling the availability of Wls for Wnt ligand export^43^. In this study, loss of Rab8a impaired Wls recycling and reduced Wnt secretion, establishing Rab8a as a regulator of the Wnt secretory pathway in the murine intestinal crypt. In parallel, the paralogue gene Rab8b can also act as a positive regulator of Wnt ligand release. Recent evidence suggests that Rab8b depletion compromises Wnt/β-catenin signalling and reduces extracellular Wnt levels *in vitro* and *in vivo*^44^. Together, these findings reveal a conserved requirement for Rab8 paralogues in Wnt trafficking and secretion. Rab8 can also attenuate Wnt/β-catenin signalling via receptor clearance. In mouse embryonic fibroblasts (MEFs), Rab8 facilitates the removal of the bona fide Wnt coreceptor Lrp6 from the plasma membrane, leading to the clearance of the active Wnt signalosome and consequently attenuating signalling^45^, suggesting a context-specific interaction of Rab8 and Wnt/β-catenin signalling. Our data in the zebrafish embryo extend these observations by showing that Rab8a does not merely facilitate Wnt export at the plasma membrane but actively drives the intracellular transport of Wnt8a in vesicles along cytonemes, delivering the ligand directly to receiving cells. Thus, previous work on Rab8a–Wls coupling and Rab8b-dependent Wnt secretion supports the functional model we uncover here: Rab8a-positive vesicles constitute a core transport module for Wnt ligands, operating in long-range cytoneme-based signalling (Extended Data 6).

### Wnt8a and Rab8a facilitate cytoneme formation

We find that overexpression of Rab8a can lead to the formation of long cytonemes, whereas inhibition of Rab8a results in fewer, shorter filopodia (Fig. 1). This observation aligns with published data suggesting that overexpression of Rab8 promotes the formation of filopodia and lamellipodia, whereas depletion of Rab8 from cells has the opposite effect^41,47^. Rab8a-positive vesicles may, therefore, have a direct structural role in cytoneme biogenesis by supplying membrane material required for protrusion elongation, particularly at the growing tip. This idea is consistent with the enrichment of Rab8a vesicles in cytoneme heads (Fig. 2) and with the Rab11a-Rab8a-VAMP3 model proposed for tunnelling nanotubes, in which Rab8a-dependent vesicles fuse with the plasma membrane to provide lipids and specific proteins required for membrane protrusion formation and growth (Zhu et al., 2018). In parallel, recent reports demonstrate that Wnt8a (and Wnt5b) activate the cytoneme formation machinery^13, 14, 46^.

These data indicate that the vesicle RABGTPase Rab8a and the cargo Wnt8a can support the generation of Wnt cytonemes and thus facilitate their long-range spreading.

### Vesicular transport in cytonemes

Studies in Drosophila proposed that morphogens are transported into cytonemes in discrete clusters, most likely in vesicles^48^. In particular, Hh was shown to associate with Rab11-positive multivesicular bodies (MVB) and the co-receptor iHog, supporting a model in which vesicular trafficking into cytonemes and the subsequent local release of exosomes underlie graded, contact-dependent signalling^49, 50^. In vertebrates, similar principles appear to apply across signalling systems. For example, telocyte-derived filopodia delivering Wnt2 in the intestinal stem cell niche display striking ultrastructural similarity to cytonemes, including a thin shaft and an enlarged, synapse-like distal domain, which we termed cytoneme head (Fig. 2)^18^. Shh cytonemes likewise transport ligand in vesicular carriers, with the actin-based motor Myo10 driving vesicle movement towards cytoneme tips and enabling rapid, contact-dependent signal activation^28^, similar to Wnt8a/Rab8a vesicles (Extended Data 3). These findings, together with our observations that Wnt8a and Wls are packaged together and subsequently co-transported in Rab8a-positive vesicles along cytonemes, suggest that cytonemes across species employ related structural principles but can use distinct vesicle types to facilitate morphogen delivery along cytonemes in specific developmental contexts.

### Implications for our understanding of morphogen function and tissue patterning

Classical models of morphogen gradient formation describe ligands as freely or hindered diffusing molecules whose spatial profiles are determined by extracellular diffusion and clearance through binding and degradation^51, 52^. In such frameworks, cell-intrinsic processes, including secretion, endocytosis, and recycling, are typically subsumed into effective parameters. Although cytoneme-mediated transport has been proposed as an alternative mechanism, it has largely remained a conceptual model, with limited direct evidence for how morphogens are physically conveyed across tissues *in vivo*. By directly visualising Wnt8a cargo within Rab8a-positive vesicles that move periodically along cytonemes and deliver ligands at discrete cell–cell contact sites (Fig. 3), our data support a model in which a substantial fraction of morphogen transport occurs intracellularly and in a vectorial manner. Consistently, there is now compelling evidence that cytonemes are employed by multiple morphogenetic signalling systems, including FGF and Dpp in Drosophila and Shh in the murine spinal cord, where they contribute to the formation, spatial distribution, and refinement of signalling gradients across developing tissues^28, 53–56^. Together, these findings suggest that the effective range and dynamics of morphogen signalling may be governed less by Brownian diffusion through extracellular space and more by regulated vesicle trafficking, motor activity, and cytoneme architecture. While extracellular diffusion undoubtedly contributes to morphogen dispersal, our results argue that morphogen gradients should be reconsidered as emergent outcomes of active, cell-controlled transport processes that spatially and temporally constrain ligand release.

## Acknowledgements

Research in the Scholpp lab, including G.S. and L.B., is supported by the BBSRC (Research Grants BB/S016295/1 and OPP490 as well as a BBSRC Equipment grant, BB/T017899/1), by the Wellcome Trust Discovery Award 8438235 and by the Living Systems Institute, University of Exeter. The Moreno-Mateos lab is supported by grants PID2024-158755NB-I00 and PID2021-127535NB-I00 funded by MICIU/AEI/ 10.13039/501100011033 and by “ERDF/EU” and Ayuda «Excelencia María de Maeztu» CEX2020-001088-funded by MICIU/AEI/ 10.13039/501100011033. The CABD is an institution funded by Pablo de Olavide University, Consejo Superior de Investigaciones Científicas (CSIC), and Junta de Andalucía. LH-H was supported by a predoctoral fellowship from the Spanish Ministry of Science and Innovation under the State Plan for Scientific and Technical Research and Innovation (2017–2020). We would further like to thank David Virshup (DUKE-NUS, Singapore), Tim Saunders (University of Warwick) and the entire Scholpp lab for their critical comments on the manuscript and the Aquatic Resource Centre and the Exter BioImaging Centre for their support.

## Materials and Methods

### Proximity ligation assay of Rab proteins in cell protrusions

The isolation procedure for cellular extension was based on previous protocols^57^. Briefly, pre-incubated 1 µg/ml of fibronectin for 2 h in 150-mm dishes, and then cells were plated at 50-60% confluence in complete medium for 12 hours. The mixture of cells and cellular extensions was collected by digestion with trypsin at a concentration of 0.125%. The supernatant was collected by centrifugation at 1000 rpm for 10 minutes and 4000 rpm for 20 minutes, and then centrifuged at 23,000 ×g for 35 minutes with a 45TI (Beckman) rotor to collect the precipitate. The precipitate was resuspended in PBS and further purified using a series of iodixanol density gradients (from 10% to 50%). The density gradients were centrifuged at 150,000 ×g for 4 h at 4°C with a SW-55Ti rotor (Beckman). Samples were collected in the layer near the 20% gradient (approximately 600µl) and mixed thoroughly with 600µl of PBS, and were centrifuged at 23,000 ×g for 35 minutes. The samples were compatible with mass spectrometry and Western blot.

### Maternal *rab8a* mRNA quantification

For RT–qPCR, total RNA was extracted from 10 embryos per biological replicate. Embryos were collected at 4 hpf, snap-frozen, and processed using the TRIzol protocol (ThermoFisher Scientific). cDNA was generated from 1000 ng RNA with the iScript kit (Bio-Rad, 1708890). qPCR reactions (10 µl) contained 2 µl of 1:5 diluted cDNA, 2 µM primers, and 5 µl of SYBR® Premix Ex Taq (Takara). Cycling conditions: 95 °C for 30 s; 40 × (95 °C for 10 s, 60 °C for 30 s). *taf15* mRNA served as the normalisation control.

### Cell Culture/Plasmids/transfection

Zebrafish PAC2 fibroblasts were cultured at 28°C without CO_2_ in Leibovitz’s L-15 Medium (Gibco, 11415064). Cells were routinely tested for mycoplasma on a three-monthly basis via PCR and yearly by broth testing. Cells were seeded on 35mm glass-bottom dishes (Thistle Scientific 81218-200), and 24 hours later, transfected using Fugene (Promega E2312) with plasmids of interest for 24-48hours. Plasmids: eGFP-Rab8a cloned into PCS2+ via ClaI/XbaI, untagged Rab8a cloned into PCS2+ via (EcoRI, XbaI), Wnt8a-mCherry in PCS2+, Wnt8a-GFP in PCS2+, Wnt5b-mCherry in PCS2+, untagged Wnt8a-mCherry in PCS2+, GFP-Rab8a (T22N) (Addgene #24899), Gap43-GFP in PCS2+, Membrane-BFP, mCherry-Myo10 (Addgene #139780) cloned into PCS2+ via Gibson, WLS-mCherry cloned into PCS2+ via ClaI/EcoRI, pEGFP-C3-Sec8 (Addgene #53758), mTurquoise2-B4GALT1(Addgene #36205) cloned into PCS2+ via Gibson and mTurquoise2 tag substituted for mCherry via (XbaI, AgeI) .

HEK293T cells were cultured in DMEM/F-12 Glutamax with 10% fetal bovine serum (FBS) at 37°C, 5% CO_2_, and 100% humidity. Cells were routinely tested for mycoplasma on a three-monthly basis via PCR and yearly by broth testing. HEK293T cells were seeded on coverslips and 24 hours later transfected with lipofectamine LTX (Thermofisher 15338100) with the following plasmids: glycophosphatidylinositol (GPI)-anchored, membrane-bound mCherry (mem –mCherry) and pcDNA-Wnt3 (Addgene #35909).

### Immunostaining of HEK2G3T cells

HEK293T cells were seeded on coverslips in six-well plates and transfected as above. After 24 h, cells were fixed in 0.25% Mem-Fix (0.1 M Sorensen’s phosphate buffer (pH 7.4), 4% formaldehyde, 0.25% glutaraldehyde) for 7 min at 4 °C^29^. Cell samples were then reduced in 0.1% Sodium Borohydride for 7 min at room temperature and then 3 x 10 min in PBS + 50nM Glycine at room temperature.

Cells were permeabilized in goat permeabilization buffer (0.1% Triton X-100, 5% goat serum, 0.2 M glycine, 1 x PBS) for 1 h at room temperature. Rab8a primary antibody (Cell signalling technology #6975) was used at 1:200 dilution in goat incubation buffer (0.1% Tween-20, 5% goat serum, 1 x PBS). 30µl of primary antibody in incubation buffer was placed on parafilm, and coverslips were placed cell-side down onto the buffer. Cells were incubated in primary antibody overnight at 4 °C. Coverslips were placed cell-side up in 6-well plates and washed 3 x 5 min in PBS. Appropriate secondary antibody (Goat anti-rabbit IgG HCL Alexa Fluor 488, ab150077, Abcam) was prepared at 1:1000 dilution in goat incubation buffer. 30µl was placed on parafilm, and coverslips were placed cell-side down onto the buffer for 1 h at room temperature. Coverslips were then placed cell-side up in 6-well plates and washed 3 x 4 min in PBS, then 1 x 3 min in MilliQ H_2_O. Coverslips were mounted on slides using ProLong Diamond (Invitrogen) and left in the dark for 24 h before imaging.

### Confocal Imaging

Cells were imaged on a Leica SP8 confocal, and embryos were imaged on a Leica SP8 Hyvolution system at 63x. Images were processed and analysed in ImageJ/FIJI.

### Super-resolution imaging

HEK293T cells were imaged on a Zeiss Elyra system with a 63 x 1.46 objective. Images were processed and analysed using Zeiss Zen software.

### 2D-CLEM workflow

Cells were cultured in Leibovitz’s L-15 medium and seeded onto gridded glass-bottom dishes. The following day, cells were transiently transfected with plasmids encoding Wnt8a-mCherry and Rab8a-GFP using Fugene (Promega). Cells were briefly fixed at 4 °C with mem-fix, counterstained with DAPI, and imaged by confocal microscopy using a Zeiss Airyscan system with a 40× 1.4NA oil-immersion objective to identify regions of interest (ROI). The imaging workflow consisted of four major steps: (1) mapping the quadrant of interest using tile scanning; (2) screening for cells of interest and recording their coordinates in ZEN Black; (3) optimising the focal plane for cytonemes and puncta, followed by acquisition of a 4 × 4 tile using transmitted light; and (4) acquiring high-resolution images using Airyscan super-resolution (SR) mode. The transmitted light tile images were used to locate regions of interest (ROIs) during TEM imaging, while the Airyscan images were used for correlation with high-magnification TEM datasets.

Samples were subsequently further fixed in aldehyde-based fixative and processed for electron microscopy. After osmium post-fixation, cells were dehydrated through a graded ethanol series and embedded in Durcupan resin. Regions corresponding to previously imaged cells were excised, polymerised, and prepared for ultramicrotomy. Ultrathin sections (70 nm) were collected on formvar-coated copper grids, contrasted with lead citrate, and imaged using a JEOL JEM-1400 transmission electron microscope operated at 120 kV.

Correlation of light and electron microscopy data was performed using the eC-CLEM plugin within the ICY bioimage analysis platform. Fluorescence images were pre-processed by channel separation and generation of maximum intensity projections prior to registration. The electron microscopy (EM) image was resampled to match the pixel size of the fluorescence microscopy (LM) dataset, and both images were cropped to the overlapping field of view. Landmark-based registration was performed by manually selecting corresponding features across modalities, and a similarity transformation (rigid with scaling) was applied, with affine refinement where necessary. The resulting transformation was then applied to the full-resolution dataset to generate high-resolution correlated overlays.

### Zebrafish maintenance and husbandry

Wildtype (WIK) zebrafish (*Danio rerio*) were maintained at 28°C on a 14-hour light/10-hour dark cycle. Experimental procedures with zebrafish were carried out in accordance with the European Communities Council Directive (2010/63/EU) and Animals and Scientific Procedures Act (ASPA) 1986. Adult zebrafish for breeding were housed and handled according to the ASPA animal care regulations. Experiments on zebrafish embryos were carried out before independent feeding at 120 hpf. All husbandry and experimental procedures were conducted under personal and project licenses granted by the UK Home Office and approved by the University of Exeter’s Animal Welfare and Ethical Review Body.

### mRNA G Cas13d/gRNA microinjection

mRNA for indicated constructs was generated using an mMessage mMachine transcription kit (Thermofisher, AM1340) from linearised plasmid. mRNA was injected at the indicated concentrations. Dextran, Fluorescein, and Biotin, 10000 MW (mini-Emerald, Invitrogen, D7178) was co-injected with mRNA to label producing cells or receiving cells. RfxCas13d purified protein was produced following a previously detailed protocol^35^. Six gRNAs were designed using the RNA-targeting tool^58^, three targeting the coding sequence (CDS) and three targeting the 3′UTR of *rab8a* mRNA. The CDS gRNAs were generated by fill-in PCR and subsequently transcribed in vitro using the HiScribe® T7 High Yield RNA Synthesis Kit, according to the manufacturer’s instructions. The resulting IVT products were purified as described previously^35^, quantified using Qubit, aliquoted, and stored at −80 °C. gRNAs targeting the 3’ UTR were purchased from Synthego Corporation (https://www.synthego.com/). A 1:1 ratio of Cas13d and a mix of 3 gRNAs targeting Rab8a was injected at 1-cell or 16-cell stage, as specified. Embryos were incubated at 28°C before mounted in low-melting agarose on 35mm plastic dishes at 50-60% epiboly for imaging or fixed in 4% PFA or analysed at 1dpf for phenotype imaging or further processing by *in situ* hybridisation.

For the genetic complementation assay, *rab8a* mRNA was obtained by PCR amplification of the CDS using specific primers and cDNA from 2.5 hpf zebrafish embryos, followed by cloning into the pT3TS vector. The resulting plasmid constructs were linearised with XbaI for 2 h at 37 °C and used as templates for *in vitro* transcription with the mMESSAGE mMACHINE™ T3 kit (#AM1348, Thermo Fisher Scientific), following the manufacturer’s protocol. Synthesised mRNA was purified using the RNeasy Mini Kit (#74104, Qiagen) and quantified with a NanoDrop spectrophotometer.

### Inhibitor treatment

Embryos were dechorionated and microinjected as described above. 1µM Wnt-C59 (TargetMol Chemicals Inc), dissolved in DMSO, was added to the media. Embryos were imaged after 3 hours of treatment.

### *In situ* hybridisation

Antisense *gbx1* and *pax6a* digoxigenin probes were generated from linearised plasmids using an RNA-labelling detection kit (Roche). Probes were purified using MicroSpin columns^14^.

*In situ* hybridisation was carried out as previously described^13^. To identify injected cell clones, following *gbx1* staining, embryos were stained with anti-FITC (product code)

### Statistical analysis

Statistics were performed and graphs generated using PRISM (Version…) and RStudio (4.4.0). Data were firstly analysed for normality and then the appropriate statistical test chosen as a result, as outlined in the figure legends. Three independent experiments were carried out unless otherwise indicated.

