## Supplementary data for "Rab8a-positive vesicles transport Wnt8a along cytonemes in zebrafish embryogenesis"

### Extended Data for Rab8a-positive vesicles transport Wnt8a along cytonemes in zebrafish embryogenesis.

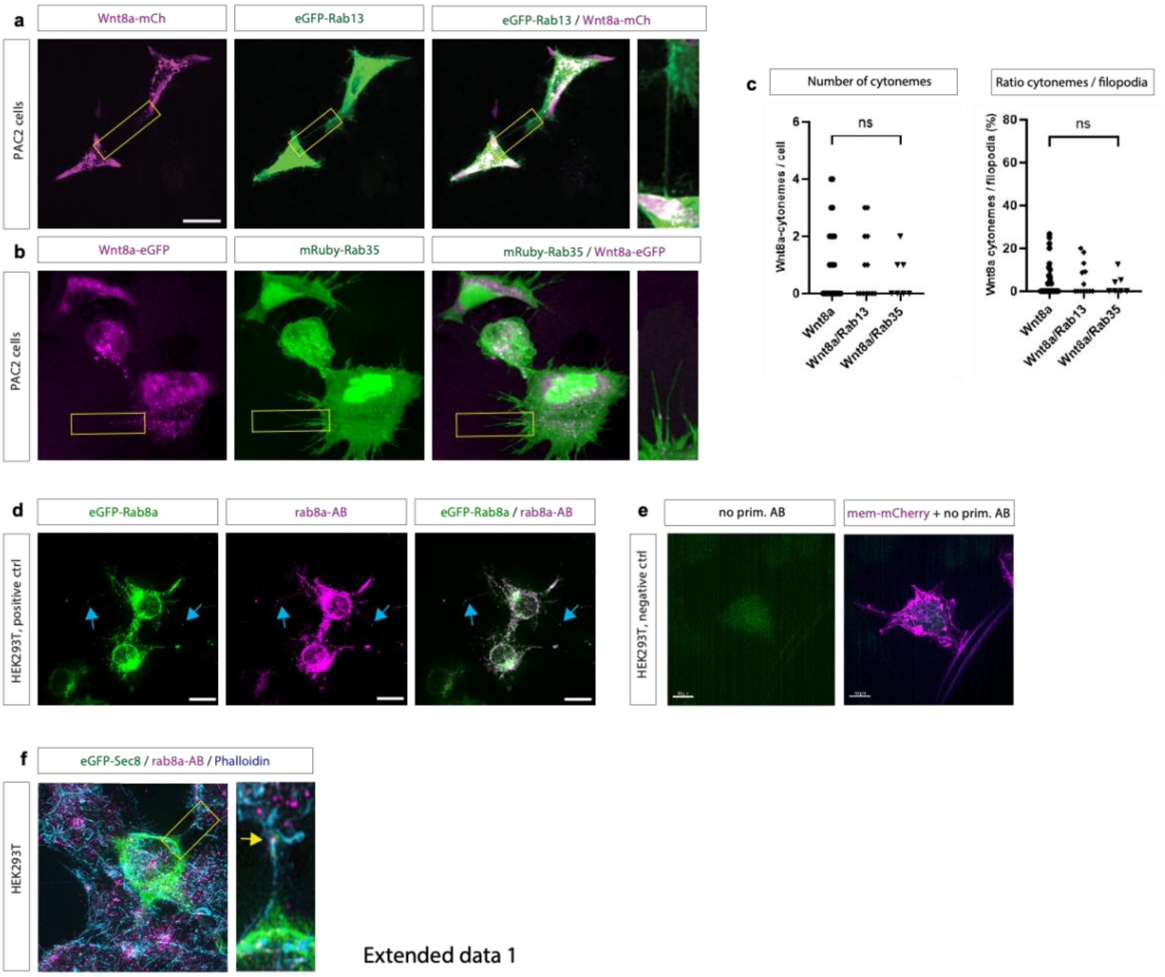


**Extended Data 1. Rab13 and Rab35 do not phenocopy Rab8a in cytoneme-associated Wnt8a output assays.**

**a,b**, PAC2 cells co-expressing Wnt8a–mCherry with eGFP–Rab13 **(a)** or Wnt8a-eGFP and mRuby-Rab35 **(b);** merged images and enlargements detect no visible co-localisation on cytonemes (boxed regions). **c,** quantification of the changes in the number of cytonemes and the ratio between cytonemes and filopodia across Wnt8a-alone, Wnt8a+Rab13, and Wnt8a+Rab35 conditions (ns as indicated). **d, e,** IHC control experiments: Rab8a antibody co-localises strongly with the expressed eGFP-Rab8a in HEK293T cells (blue arrows), whereas no signal was detected when the primary antibody was lacking. **f,** IHC analysis of HEK293T cells transfected with eGFP-Sec8 and stained with Phalloidin-405 reveals co-localisation of endogenous Rab8a puncta with the post-Golgi marker on cytoneme shafts (yellow arrows).

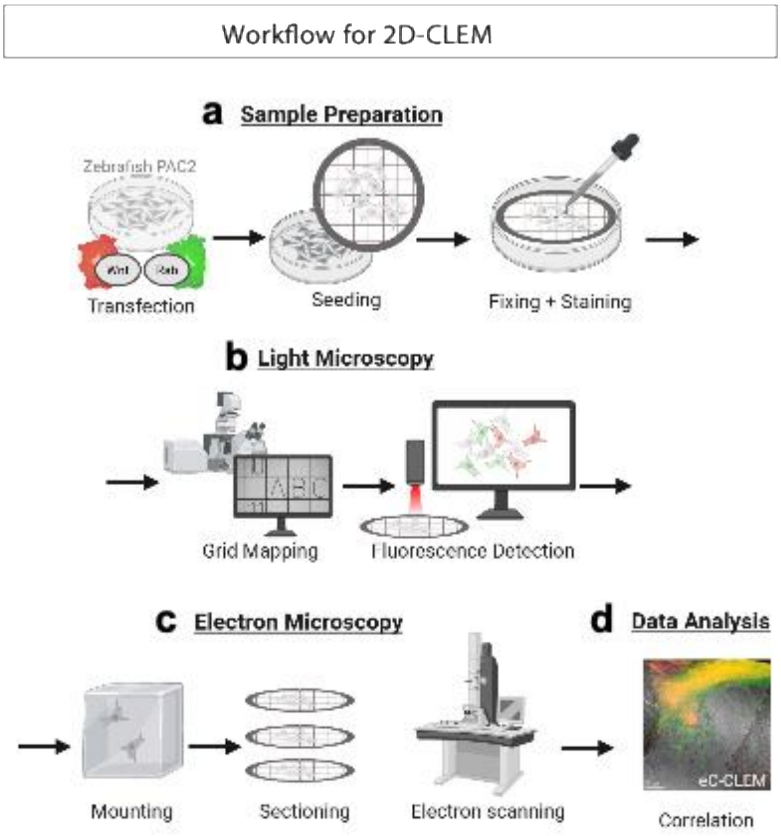


**Extended Data 2. Workflow for 2D correlative light and electron microscopy (2D-CLEM)**
Schematic of the 2D-CLEM pipeline: **a,** transfection of PAC2 cells, seeding on gridded supports, fixation and staining; **b,** grid mapping and fluorescence imaging; **c,** mounting, sectioning and electron microscopy; **d,** image registration and correlation for eC-CLEM analysis.

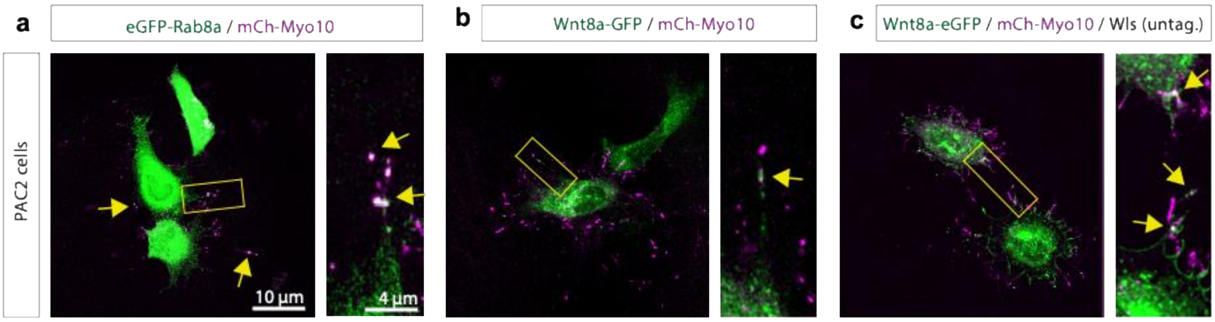


**Extended Data 3. Myo10-labelled cytonemes carry Rab8a/Wnt8a puncta *in vitro***
**a–c,** PAC2 cells expressing mCherry-Myo10 with eGFP–Rab8a (a), Wnt8a–GFP (b), or Wnt8a–eGFP together with mCherry-Myo10 and untagged Wls (c). Insets show boxed regions; arrows mark puncta along Myo10-positive cytonemes. Scale bars as indicated.

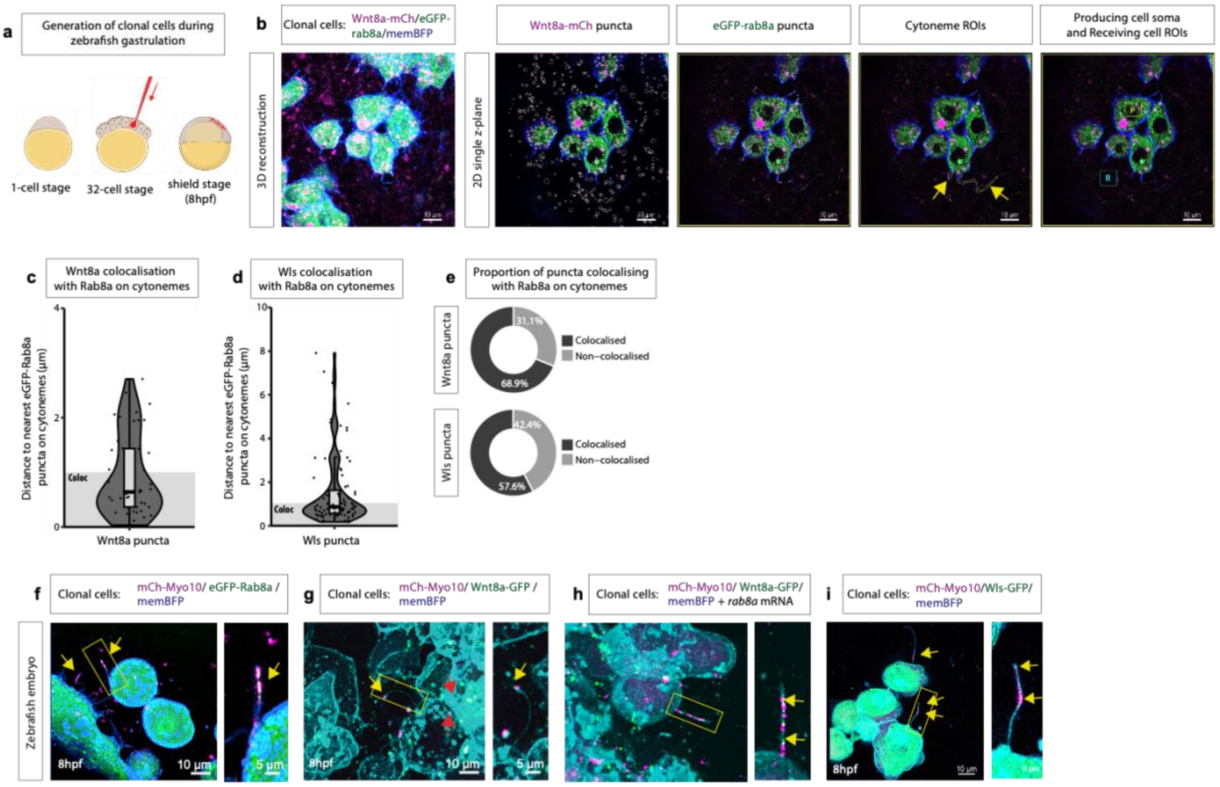


**Extended Data 4. Quantitative analysis of cytonemes in the zebrafish gastrula carrying Rab8a/Wnt8a puncta**

**a**, Schematic illustrating mosaic clone generation: injection at the 32-cell stage to label sparse clonal populations analysed at shield stage (8 hpf). **b**, Zebrafish embryos (8hpf) with clonal expression of Wnt8a-mCherry, eGFP-rab8a and memBFP. 2D single z-plane panels illustrating how Wnt8a-mCherry and eGFP-rab8a puncta are isolated for analysis. Arrows indicate selected ROIs representing cytonemes. Boxed regions show examples of ROIs within the producing cell soma (P) and in receiving cells surrounding the clone (R).**c,d,** quantification of the distance of cytonemal Wnt8a-mCherry puncta (**c**) or Wls-mCherry puncta (**d**) from the nearest eGFP-rab8a puncta. Distances of <1µm are classified as co-localised. **e,** Proportion of cytonemal Wnt8a-mCherry and Wls-mCherry puncta which colocalise withe eGFP-rab8a. **f-i,** Zebrafish embryos (8 hpf) with clonal expression of mCherry-Myo10, eGFP-rab8a and memBFP (**j**),Wnt8a–GFP, mCherry-Myo10 and memBFP (**k**) or with additional *rab8a* mRNA (**h**), and mCherry-Myo10, Wls-eGFP and memBFP (**i**); insets show boxed regions and puncta along protrusions. Arrows indicate co-localisation of fluorescent puncta on cytonemes. Scale bars as indicated.

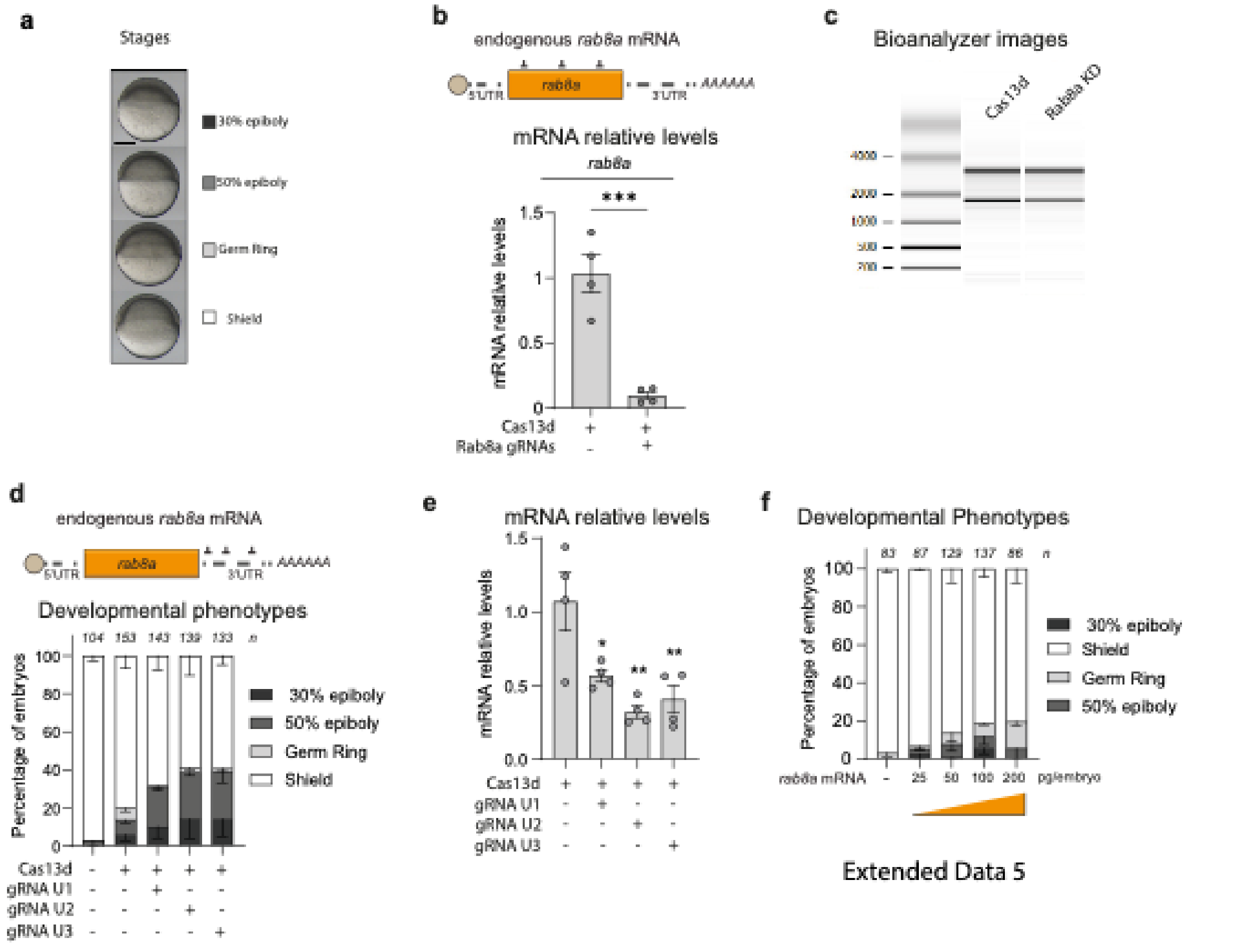


**Extended Data 5. RfxCas13d-based knockdown of Rab8a in zebrafish.**

**a,** Reference images defining developmental staging categories used for phenotypic scoring (30% epiboly, 50% epiboly, germ ring, shield). **b,** Schematic representation of the *rab8a* mRNA and the gRNAs used to target the coding sequence (CDS). The 5'UTR, 3'UTR, and poly(A) sequence are indicated. Bottom. Relative *rab8a* mRNA levels under the indicated conditions, analysed by RT-qPCR. Results are shown as the mean ± standard error of the mean from two experiments, each including two biological replicates (n = 10 embryos per biological replicate). *taf15* mRNA was used as the normalization control. (****p* < 0.001, unpaired t-test). **c,** RNA integrity profiles analysed with the Agilent 2100 Bioanalyzer for RNA samples collected at 6 hpf from pools of 10 embryos injected only with RfxCas13d protein (Cas13d) or with RfxCas13d together with a mix of three gRNAs targeting *rab8a* mRNA (Rab8a KD). **d,** Top. Schematic representation of the *rab8a* mRNA and the gRNAs used to target its 3'UTR. The 5'UTR, 3'UTR, and poly(A) sequence are indicated. Bottom. Stacked bar plots showing the percentage of embryos observed at 6 hpf for control embryos (uninjected), embryos injected only with RfxCas13d protein, and embryos injected RfxCas13d protein with different individual gRNAs targeting the 3'UTR of *rab8a* mRNA. The results represent the mean of three independent experiments ± the standard error for each developmental stage (**a**). **e,** Relative *rab8a* mRNA levels under the indicated conditions, analyzed by RT-qPCR. Results are presented as the mean ± standard error of the mean from two experiments, each with two biological replicates (n = 10 embryos per biological replicate). *taf15* mRNA was used as the normalization control. (ns = non-significant, *p < 0.05, **p < 0.01, one-way ANOVA). **f,** Stacked bar plots showing the percentage of embryos observed at 6 hpf for control embryos (uninjected) or embryos injected with increasing amounts of *rab8a* mRNA. The results represent the mean of two independent experiments ± the standard error for each developmental stage (**a**).

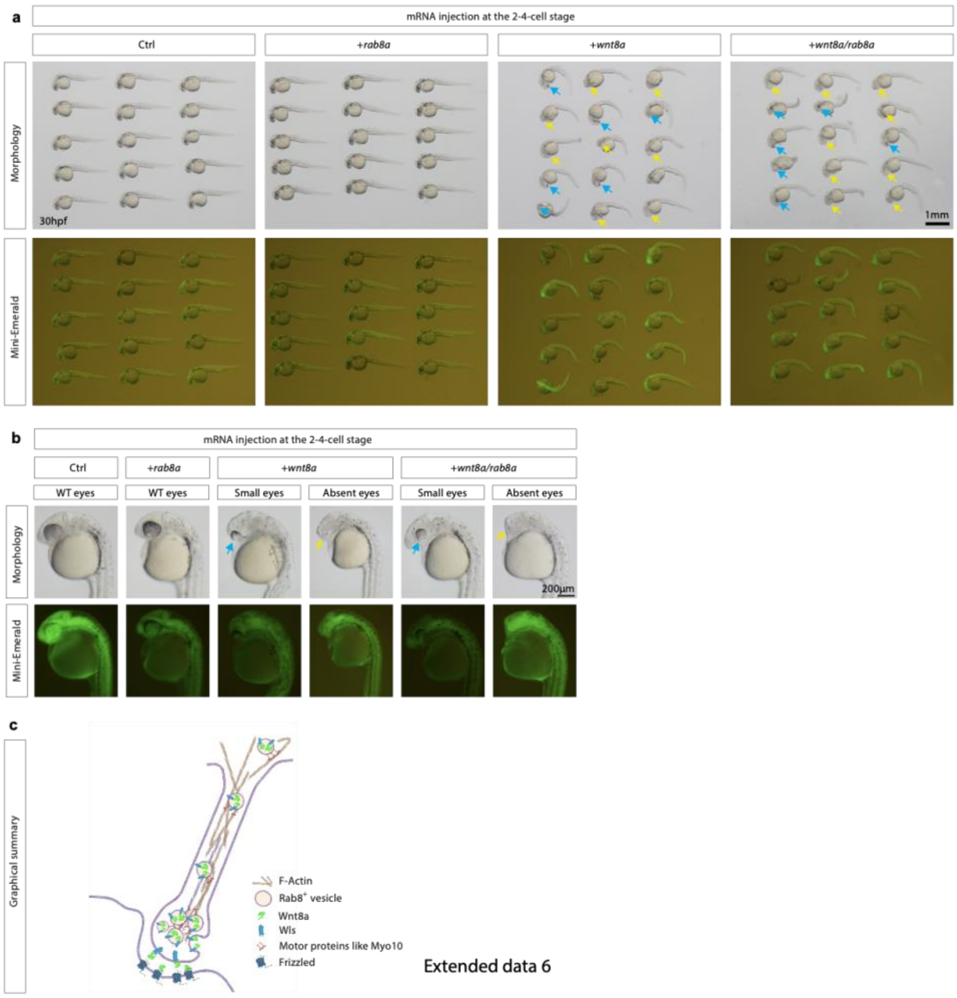


**Extended Data 6. Rab8a co-expression modulates Wnt8a overexpression phenotypes**

**a,** Representative embryos at 30 hpf after mRNA injection at the 2–4 cell stage (Ctrl, +rab8a, +wnt8a, +wnt8a/rab8a) shown in brightfield (top) and with lineage tracer fluorescence (Mini-Emerald; bottom); arrows indicate characteristic Wnt8a-associated morphological abnormalities. **b,** Higher-magnification views highlighting eye phenotypes (WT, small, absent) under the indicated injection conditions, with corresponding Mini-Emerald fluorescence. Scale bars as indicated. **c,** Graphical model summarising cytoneme-based, Rab8a-vesicular transport of Wnt8a together with Wls toward cytoneme tips and delivery to Frizzled-positive receiving cells.
